# Deep neural network and field experiments reveal how transparent wing windows reduce detectability in moths

**DOI:** 10.1101/2020.11.27.401497

**Authors:** Mónica Arias, Cynthia Tedore, Marianne Elias, Lucie Leroy, Clément Madec, Louane Matos, Julien P. Renoult, Doris Gomez

**Affiliations:** CEFE, CNRS, Univ. Montpellier, Univ. Paul Valéry Montpellier 3, EPHE, IRD, Montpellier, France; ISYEB, CNRS, MNHN, Sorbonne Univ., EPHE, Univ. Antilles, 45 rue Buffon CP50, Paris, France; Univ. Hamburg, Faculty of Mathematics, Informatics and Natural Sciences, Institute of Zoology, Hamburg, Germany; INSP, CNRS, Sorbonne Univ., Paris, France

**Keywords:** background matching, disruptive coloration, transparency, wild bird predators, artificial prey, deep learning

## Abstract

Lepidoptera – a group of insects in which wing transparency has arisen multiple times - exhibit much variation in the size and position of transparent wing zones. However, little is known as to how this variability affects detectability. Here, we test how the size and position of transparent elements affect predation of artificial moths by wild birds in the field. We also test whether deep neural networks (DNNs) might be a reasonable proxy for live predators, as this would enable one to rapidly test a larger range of hypotheses than is possible with live animals. We compare our field results with results from six different DNN architectures (AlexNet, VGG-16, VGG-19, ResNet-18, SqueezeNet, and GoogLeNet). Our field experiment demonstrated the effectiveness of transparent elements touching wing borders at reducing detectability, but showed no effect of transparent element size. DNN simulations only partly matched field results, as larger transparent elements were also harder for DNNs to detect. The lack of consistency between wild predators’ and DNNs’ responses raises questions about what both experiments were effectively testing, what is perceived by each predator type, and whether DNNs can be considered to be effective models for testing hypotheses about animal perception and cognition.

## Introduction

The coevolutionary arms race between prey and predators has generated some of the most striking adaptations in the living world, including lures, mimicry and camouflage in prey, and highly sophisticated detection systems in predators [1]. Transparency, by definition, constitutes the perfect background matching against virtually all types of backgrounds. Transparency is common in pelagic environments where there is no place to hide [2], and often concerns the entire body because: 1) small differences in refractive index between water and biological tissues limit reflections that would betray the presence of a prey, 2) ultraviolet-protective pigments are not necessary in the depths of oceans because UV radiation is filtered out by water and, 3) water provides support, limiting the need for thick, sustaining (often opaque) tissues. Contrary to aquatic species, transparency on land has only evolved as “elements” (i.e. parts of their body). It is currently unknown which visual configurations of transparent and opaque elements are the best at reducing prey detectability on land.

Unlike in aquatic taxa, transparency has evolved in few terrestrial groups, and is mostly restricted to insect wings. Within the Lepidoptera, however, transparency has evolved several times independently in typically opaque-winged butterflies and moths. According to recent experimental evidence, the presence of well-defined transparent windows reduces detectability in both conspicuous [3] and cryptic [4] terrestrial prey. However, wing transparency can occur in different forms (Figure 1): as small windows (e.g., as in the moths *Attacus atlas,* or *Carriola ecnomoda)* or as large windows (e.g., as in the sphinx *Hemaris fuciformis)* delimited by brownish elements or conspicuous wing borders (e.g., as in the Ithomiine tribe). Wing transparency can also be associated with discontinuous borders, touching prey edges (e.g., as in the elytra of the tortoise beetle *Aspidomorpha miliaris* or in moths of the genus *Bertholdia).* Whether the size of transparent windows or their disrupting contour have any effect on prey detectability remains to be studied.

**Figure 1.**
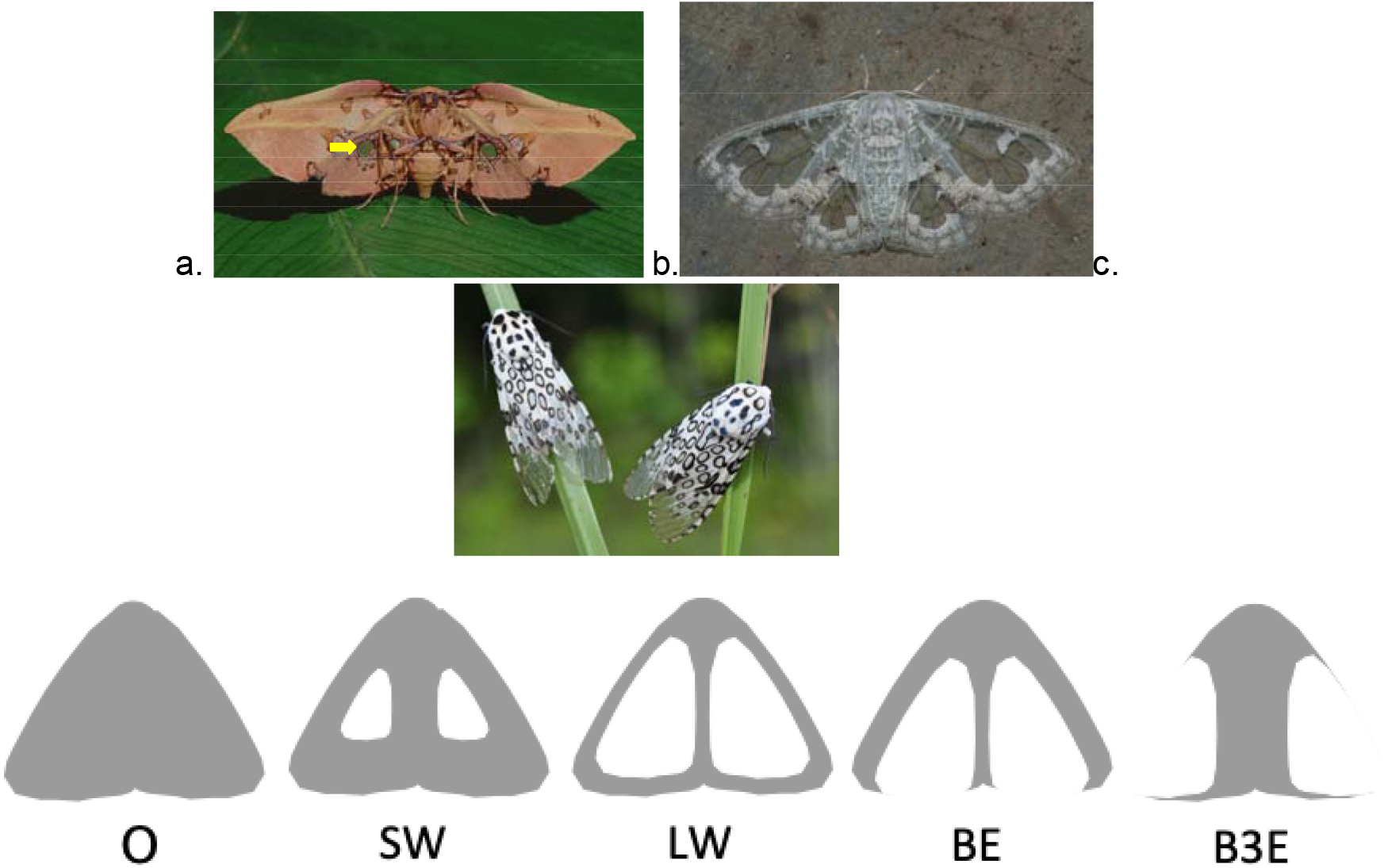
Examples of diversity of position and size of transparent elements in butterflies. a. *Siculodes aurorula* shows small windows (indicated by the yellow arrow, photo: © Adrian Hoskins), b. *Carriola thyridophora* shows large windows (photo: © Vijay Anand Ismavel), c. *Hypercompe scribonia* shows windows breaking the bottom edge (photo: © Green Futures). In the second row, 5 morphs of artificial moths used in this study.

This diversity of forms suggests that including transparent elements can reduce detectability by different mechanisms. For example, a transparent window surrounded by an opaque border could reduce detectability by increasing background resemblance. The larger the transparent area, the larger proportion of the surface area of the prey that matches its background (“background matching hypothesis”). On the other hand, transparent elements breaking the prey outline can hamper detection as a form of disruptive coloration [5,6]. In disruptively coloured prey, colours that imitate the background are combined with contrasting elements that are located on the prey border, breaking its visual edges and making it difficult to detect (“disruptive coloration hypothesis”). Broken edges have been shown to work better than background matching [5], especially when low and intermediate colour contrast elements are included [7]. Additionally, transparent elements touching prey borders can make the prey appear to be a smaller size than it actually is, reducing its perceived profitability and its risk of being attacked, as predation rate is directly correlated with prey size [8]. To understand the relative importance of these different mechanisms in partially transparent terrestrial prey, we need to investigate the effect of size and position of transparent elements.

Field experiments with artificial prey models are a common way to explore the behaviour of natural predators in response to the visual characteristics of prey and thus to infer the perceptual mechanisms of predation avoidance. This approach has been used, for instance, to reveal the defence advantages of disruptive coloration [5], which is more efficient when less symmetrical [9] and in the absence of marks attracting predators’ attention to non-vital body parts [10]. However, field experiments are time-consuming and labor-intensive (from 11 to 20 weeks per experiment in the field to obtain attack rates ranging from 11.3% to 18% [11–13]), which reduces the combination of conditions that can be tested simultaneously.

Artificial intelligence algorithms, and deep neural networks (DNNs) in particular, are becoming common tools for tackling questions in ecology and evolution while reducing human labour [14,15]. DNNs are able to represent complex stimuli, such as animal phenotypes and their colour patterns, in a high-dimensional space in which Euclidean distances can be calculated [16–18]. As in a colour space, the distance between phenotypes or patterns in the representational space of a DNN have been shown to be correlated with their perceived similarity [19].

DNNs are made up of multiple layers of neurons that lack interconnections within layers but are connected to neurons in adjacent layers. The firing intensity of a neuron depends on the strength of signal received by connected upstream neurons and the neuron’s nonlinear activation function. Learning comes about by iteratively tweaking the weight of each neuron’s output to downstream neurons until a more accurate final output is attained. Although the architecture and computational mechanics of artificial neural networks were originally inspired by biological neural networks, the primary goal of most DNNs today is to produce accurate outputs (in our case, categorizations) rather than to mimic the mechanics of biological neural networks. Indeed, this prioritization of functionality over mechanistic mimicry is true for the DNNs used in the present study. Interestingly, however, the spatial distribution of different classes of objects in the representational space of DNNs has been shown in various studies to be correlated with that of the primate visual cortex [20– 25]. This is suggestive of some degree of mechanistic similarity in the way that artificial and biological neural networks encode and categorize objects. Beside primates, the ability of DNNs to predict the responses of animals remains almost untested (but see [26]).

Here, we explored the perceptual mechanisms by which transparent wing windows decrease predation, using wild avian predators and DNNs as observers of artificial moths. As both types of observers were naïve to transparent butterflies and moths, our study also shed light on the selective advantages that could favour the evolution of these mechanisms. First, we carried out field experiments to analyse the effect of different visual characteristics of transparent windows and adjacent elements on the attack of prey by avian predators. We predicted that larger transparent windows touching prey outlines would make prey more difficult to detect, presumably *via* both background matching and disruptive coloration. Second, we studied the ability of DNNs to detect moths with varying sizes and configurations of transparent elements, aiming to compare DNN responses to the behaviour of real predators. Finally, using DNNs, we explored which characteristics of transparent elements might promote the establishment of new cryptic morphs in a given population. Our study sheds light on how well predator behaviour can be predicted by artificial intelligence.

## Material and methods

### Field experiments

We followed an experimental design similar to that described in [4]. Briefly, we performed predation experiments in April and May 2019 in two localities in southern France: La Rouvière forest (43.65°N, 3.64°E) and the Montpellier zoo (43.64°N, 3.87°E), for three 1-week sessions at each place. We monitored artificial prey survival from predation by bird local communities once per day for the four consecutive days after placing them on trunks, and removed them afterwards. Artificial prey (body and wings) were pinned on green oak *(Quercus ilex)* tree trunks (>10cm in diameter, with little or no moss cover) every 10m. For further details see “Extended Materials and Methods” in ESM.

### Artificial moths

As described by Arias *et al* [4] and similarly to other experiments, artificial moths consisted of paper wings and an edible body [5,6]. Wings consisted of right triangles resembling resting generic moths (i.e., not representing any real species). Triangle dimensions were 25mm height by 36mm width. Artificial prey had grey elements bearing low chromatic and achromatic contrasts with the oak trunk for both UV- and V-vision birds. This colour thus enabled us to explore the effect of transparent elements on reducing detectability of cryptic prey. For details on the production of these artificial moths see “Extended material and methods” in ESM.

We analysed five types of artificial grey moths with different wing characteristics (Figure 1): opaque (O morph), with small transparent windows within the wing (SW morph), with large transparent windows within the wing (LW morph), with large transparent windows touching the bottom edge of the wing (BE morph) and with large transparent windows touching all three wing edges (as each window is touching two edges of the three moth edges, B3E morph). Morphs that included transparent elements were built by cutting two right triangular windows out of the laminated grey triangle. Dimensions for each triangular window were 7 mm width by 10 mm height (transparency occupying thus 15% of original grey surface) for the SW morph, and 14mm width by 18mm height for the LW, BE and B3E morphs (transparency occupies thus 46.6% of originally grey surface). To simulate transparent wing surfaces, we then added a transparent film (3M for inkjet, chosen for its high transparency even in the UV range see ESM, Figure S1) underneath the remaining parts. On top of the moth wings, we added an artificial body made from pastry dough (428g flour, 250g lard, and 36g water, following [27]), dyed grey by mixing yellow, red and blue food dyes (spectrum in Figure S1). This malleable mixture allowed us to make small bodies on which we could record marks made by bird beaks and distinguish them from those made by insects.

### Data collection and analysis

During monitoring, we considered artificial moths as attacked by birds when their body showed V-shaped or U-shaped marks, or when the body was missing without signs of invertebrate attacks (i.e. no body scraps left on wings or around the moth on the trunk). We removed all remains of artificial moths attacked by birds, and replaced them when attacked by invertebrates or when the entire artificial prey (wings, body and pin) were missing, as we could not exclude that the prey item fell down or was blown away by the wind. Non-attacked prey were treated as censored data in the analyses (i.e. prey that survived at least until the end of the experiment). We analysed prey survival, fitting three independent Cox proportional hazard regressions [28], using as explanatory variables one of the three different ways to describe the visual characteristics of morphs: 1) per morph (O, SW, LW, BE, B3E), 2) per border characteristics (having a full wing border (O, SW, LW) or a border broken by transparent windows in one (BE) or three edges (B3E)), and 3) per transparent surface (relative size of both transparent windows (0 for O, 0.14 for SW and 0.57 for LW, BE and B3E)). We also included week as an explanatory variable in order to control for time in the season as well as place, as the experiment was never run simultaneously at La Rouviére forest and at the zoo during the same week. Interactions between morph or its characteristics and week were also included as explanatory variables in order to explore whether differences in locality and/or time in the breeding season might affect predation on different morphs. Overall significance was measured using a Wald test. Statistical analyses were performed in R [29] using the *survival* package [30]. Using the Cox survival model results, we performed pairwise post-hoc comparisons between survival of the different morphs and their characteristics. Additionally, using the Cox survival model coefficients, we produced “distance matrices” of pairwise differences of survival between morphs (survival coefficient for morph A - survival coefficient for morph B): 1) per morph, 2) per transparent surface size and 3) per border characteristics (Tables S5).

Differences in attack rates between places and/or weeks could be produced by differences in predation pressure, which can be higher when more nests are occupied and/or when more chicks at their developmental peak are present in the population. Therefore, we explored demographic effects of great and blue tit local populations on attack distribution. For further details see “Extended materials and methods” in ESM.

### Deep Neural Networks

We used deep neural networks (DNNs) for the second and third aims of this study. We trained DNNs to detect moths on tree trunks and scored their performance at detecting different moth forms. We assessed the performance of different DNNs at detecting moths, and compared the responses of DNNs and avian predators when detecting the same prey. Additionally, using DNNs, we predicted the characteristics that will render a prey morph highly cryptic and thus potentially able to establish in a population.

### Prey used

We took pictures of artificial grey moths identical to those used in the field experiments. A tablet (iPad mini 2) and a smartphone (Moto G5, Android 8.1.0) using the “U camera” application to produce square-shaped photos were used to take pictures of artificial prey. No HDR option was activated on either of the cameras. Each device was mounted on a tripod to take pictures of each of five artificial moth morphs, each pinned to the same position on a tree trunk, as well as the tree trunk alone with no moth present. This controlled setup ensured that the DNNs used distinguishing characteristics of the different morphs, rather than finding unintended correlations between each morph and other characteristics of the visual background. Each morph was photographed against 222 unique visual backgrounds.

### Selected DNNs and general training

Rather than training DNNs from scratch (i.e. initializing all neuronal outputs with random weights), we used DNNs pre-trained to correctly identify 1000 object categories from more than a million images from the ImageNet database. Pre-trained networks have already learned to detect image features common to many classes of objects, and have pre-defined neuronal weights that are useful for learning new combinations of features. We took such DNNs as a starting point, and then used transfer learning to train them on the new task of detecting artificial moths. Transfer learning replaces the last few layers of an existing DNN with new learning and classification layers that rapidly adapt to a new classification task after training on only a limited dataset. To test the robustness of results obtained from DNNs, we used six different DNNs that were similarly trained: AlexNet, VGG-16, VGG-19, ResNet-18, SqueezeNet, and GoogLeNet. Weights were frozen in the first ten layers to preserve simple feature detectors already learned from the ImageNet database. For further details about initial learning rate and solvers see “Extended material and methods” in ESM. To avoid overfitting DNNs to the training set, training was halted when the loss (a measurement of DNN uncertainty) on the validation set failed to decrease three times. At this point, training was considered complete. After training and testing, each DNN returned a score between 0 and 1 indicating its level of confidence that there was a moth in each picture. This score was determined by a sigmoid binary classifier (softmax layer in the DNN). Images with scores greater than 0.5 were classified as displaying a moth. To verify that DNNs were using features of the wings, and not the body alone, to detect moths, we generated class activation maps of the final layer of the DNNs. These maps indicated which parts of the image contributed the most towards determining that the image contained a moth. SqueezeNet, ResNet-18, and GoogLeNet can produce such maps, but AlexNet, VGG-16, and VGG-19 cannot due to the fact that their final layer consists of multiple interconnected layers. All training and testing of DNNs and generation of class activation maps was conducted using MATLAB [31].

We used two training methods to explore two different questions: 1) In a community of different moth morphs, all of which are familiar to predators, which morphs will be the most successful at evading predation? and, 2) If predators are only experienced with opaque morphs, which novel morph(s) with transparent elements will most easily establish themselves in the population? To see the distribution of images in training, validation and test set for both experiments and further details, see Figure 3 and “Extended material and methods” in ESM.

### Analyses of DNNs results

First we compared the DNNs’ performances at moth detection under different training scenarios by comparing their false positive rates (proportion of background images with scores higher than 0.5). Additionally, and for each training method, we computed summary statistics (mean and standard deviation) of DNN scores per morph and DNN architecture. As the DNNs’ performances were rather similar (Table S2), we pooled them together. We created a binomial response variable that was set to one for DNN scores higher than 0.5 and set to zero for DNN scores lower than 0.5. We fitted a Generalised Linear Mixed model assuming a binomial distribution, using a binary response variable describing whether a moth was detected (1) or not (0) in each image. The explanatory variable was the moth morph and had five levels: O, SW, LW, BE and B3E. Independent models were fitted for the two different experiments testing the contrast: a) O was more detected than other morphs, b) morphs with small windows (SW) were more detected than morphs with large windows (LW, BE and B3E), c) BE was more detected than B3E and d) morphs with complete windows (SW and LW) were more detected than morphs with broken edges (BE and B3E). Random effects included image ID, DNN architecture and replicate of each DNN architecture (100 replicates for each of the six DNN architectures). Finally, and similar to the field experiment, we produced “distance matrices” for detection differences between prey types: 1) per morph, 2) per transparent surface size and 3) per border (Tables S5). Distances were calculated as the difference between DNNs scores per morph. Applying a Mantel test, we compared results from the field and from both DNNs experiments.

## Results

### Effect of window size and position on prey survival in the field

555 artificial moths were attacked out of the 1583 used in the field experiment (35.1% of attack rate). We detected no significant differences in attacks between morphs when separating all the morphs (Wald test = 3.92, df = 4 *p =* 0.4, Figure 2). We did, however, find differences when grouping morphs according to the characteristics of their transparent windows. When transparent windows touched the border (morphs BE and B3E), prey were less attacked (*z* = −2.03, *p =* 0.04), suggesting that this characteristic renders morphs less detectable. However, predation of morphs sporting only different transparent window sizes was rather similar (morph SW vs morphs LW, BE, B3E: *z* = −1.49, *p* = 0.13). No difference in attacks was registered between the two locations (z = −2.03, *p* = 0.73). However, more attacks were registered at the zoo at the beginning of the experiment (attacks: week 2 > week 5, *z* = 4.41, *p* <0.001), and at La Rouviére forest at the end of the experiment (attacks: week 6 > week 4, *z* = 2.75, *p* = 0.006). We did not find any correlation between attacks per day or per week and predator demographic dynamics that could either increase or decrease predation pressure (the number of active nests, the number of chicks between 9 and 15 years old or the number of hatchings for great and blue tits (Figure S2, best model includes none of the demographic variables to explain the variation in attack rate).

**Figure 2.**
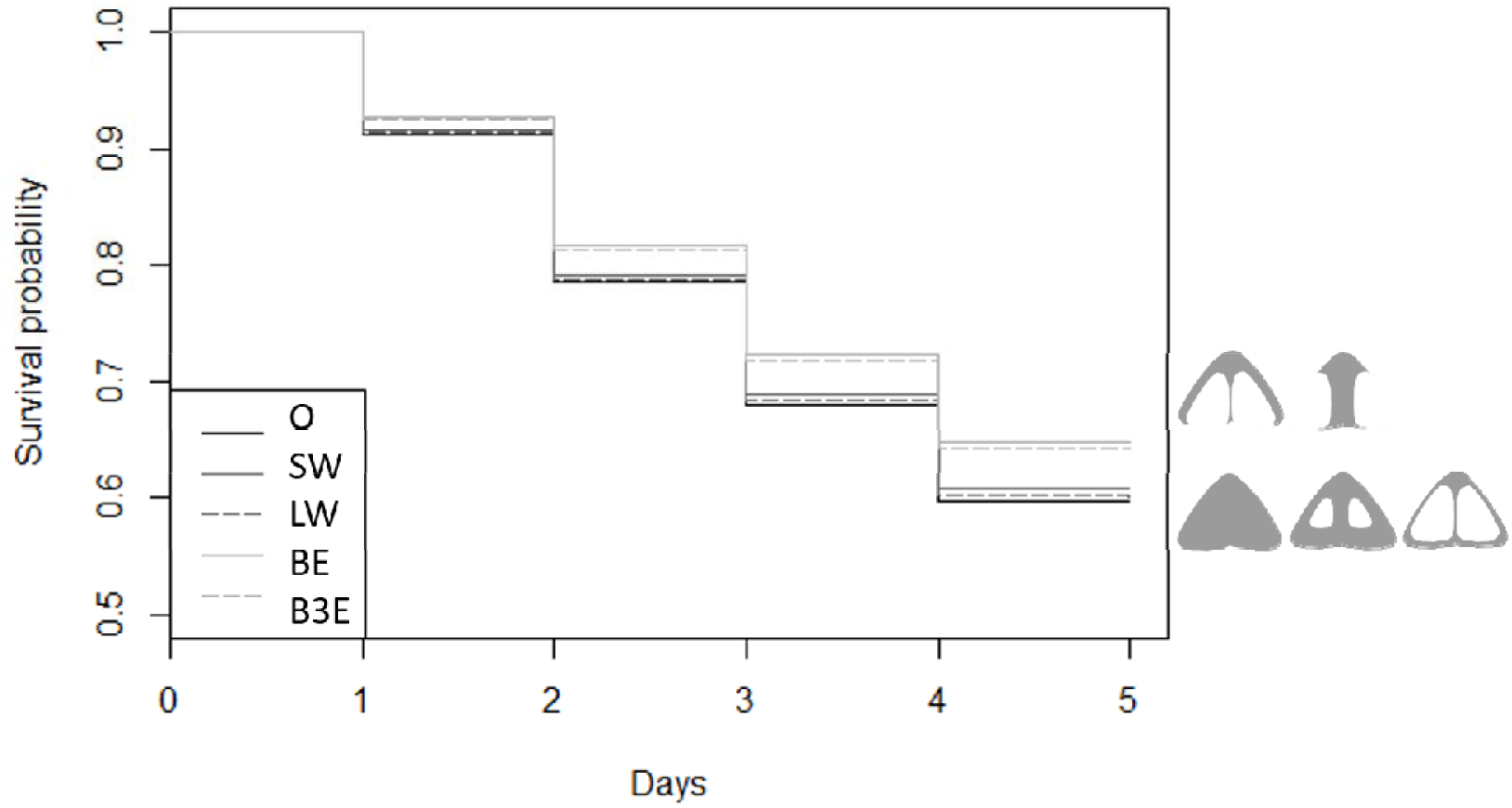
Survival of artificial prey without transparent elements (O-opaque), with small (SW) and large transparent elements touching none (LW), one (B1E) or three prey edges (B3E). Artificial butterflies were placed on tree trunks and monitored for their ‘survival’ every day for 4 days. Data from the six weeks during which the experiment was conducted are pooled together.

**Figure 3.**
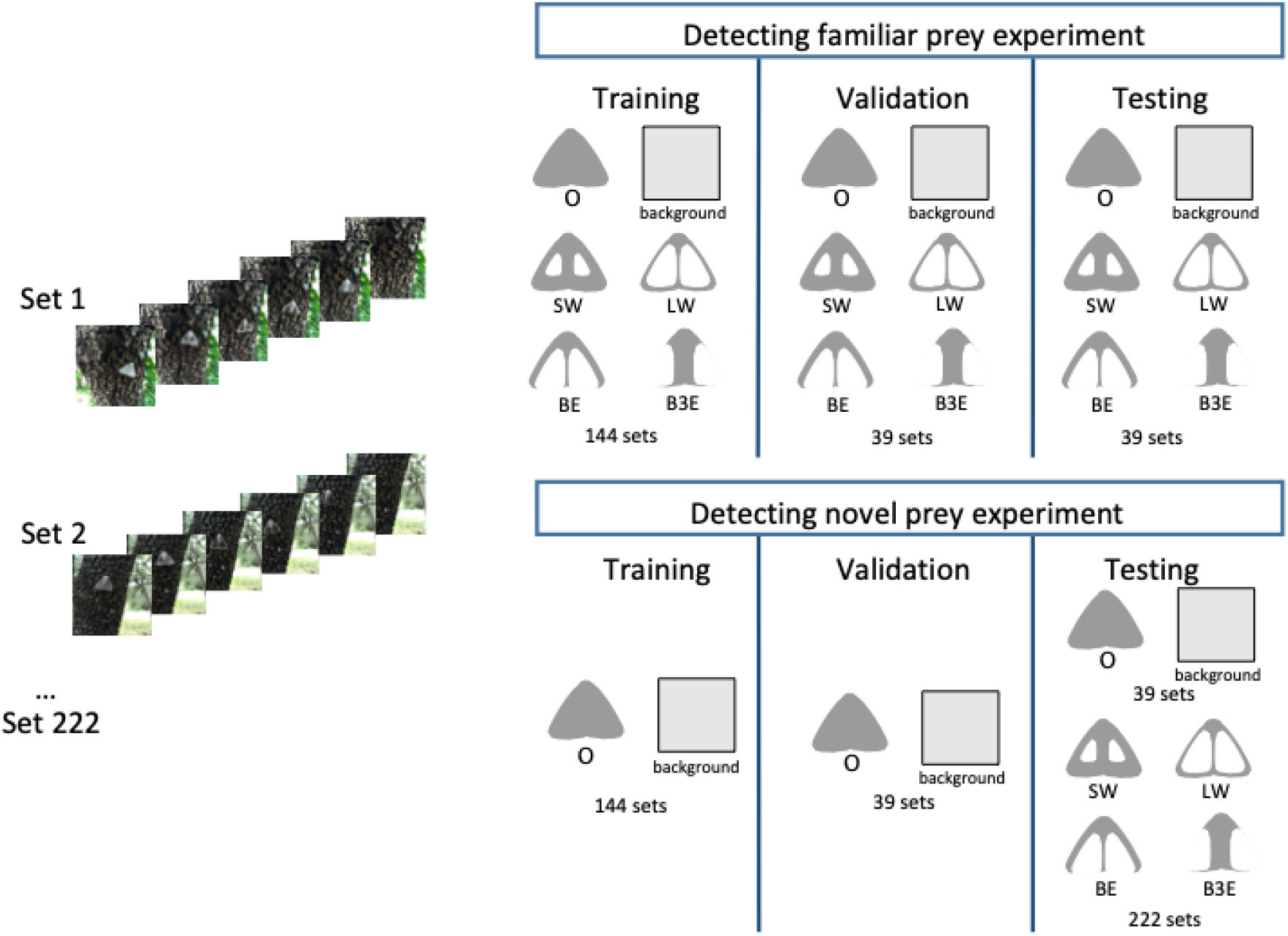
Scheme of training and testing when detecting familiar (top row) or novel prey (bottom row) with the different DNNs. Each cycle was repeated 100 times.

### Image elements used by DNNs to detect moths

Class activation maps (CAMs) showed that DNNs used features of both the wings and body to recognize moths. SqueezeNet produces the highest-resolution CAMs, so we show examples of CAMs generated by SqueezeNet for each of five morphs against two different backgrounds in Figure 4.

**Figure 4.**
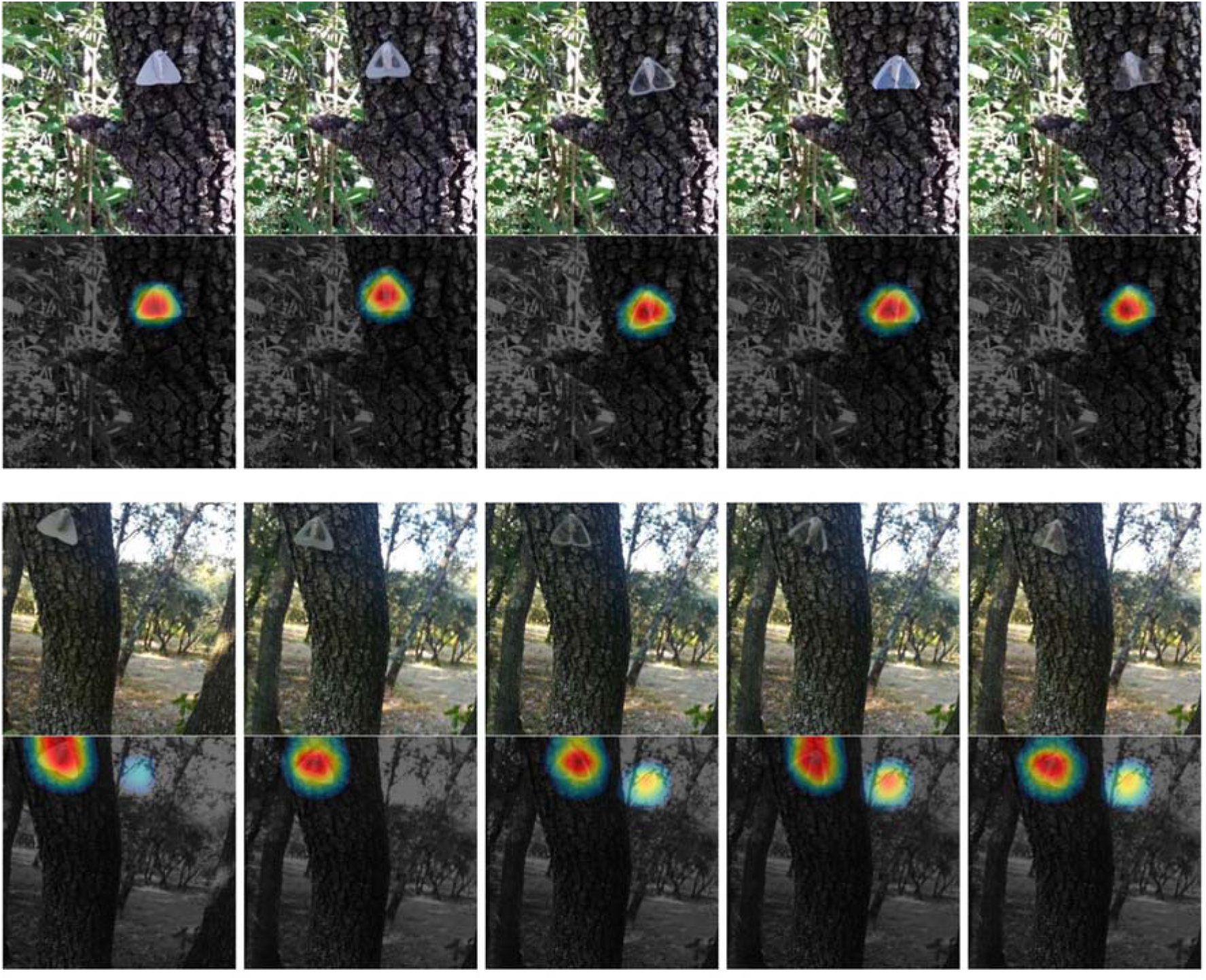
Example images of each of the five morphs against two different backgrounds, together with their corresponding class activation maps (CAMs) produced by SqueezeNet. The CAMs are heatmaps with “hot”, or red, regions corresponding to parts of the image with the highest activations.

### DNNs’ detection of moth morphs used in the field experiment

When trained only on the “O” form, AlexNet produced the highest false positive rate (0.19). The lowest was produced by ResNet (0.01). When trained on all morphs, AlexNet again produced the highest false positive rate (0.31), while GoogLeNet produced the lowest (0.09, Table S1).

Different DNN architectures and training methods produced similar results (Table S2). When pooling together all DNNs, we found that “O” was the easiest morph to detect in both experiments (detecting familiar prey experiment: z = 9.8, p<0.001; detecting novel prey experiment: z = 20.25, p<0.001, Figure 5 and Table S3). Having large windows, independently of their position (pooling together LW, BE and B3E), reduced prey detectability more than having small windows (detecting familiar prey experiment: z = 4.66, p<0.001; detecting novel prey experiment: z = 10.87, p<0.001, Table S3). Morphs having windows touching three edges were more difficult to detect than morphs touching only one edge especially when detecting familiar prey (detecting familiar prey experiment: z = 5.67, p<0.001; detecting novel pre experiment: z = 2.39, p=0.02). Finally, morphs having complete windows (pooling together SW and LW) were similarly detected than those with the broken edges (pooling together BE and B3E, detecting familiar prey experiment: z = −0.32, p=0.75, 83% detected for complete windows vs 80% for broken edges; detecting novel prey experiment: z = 0.85, p=0.4, 62.3% detected for complete windows vs 48.3% detected for broken edges).

**Figure 5.**
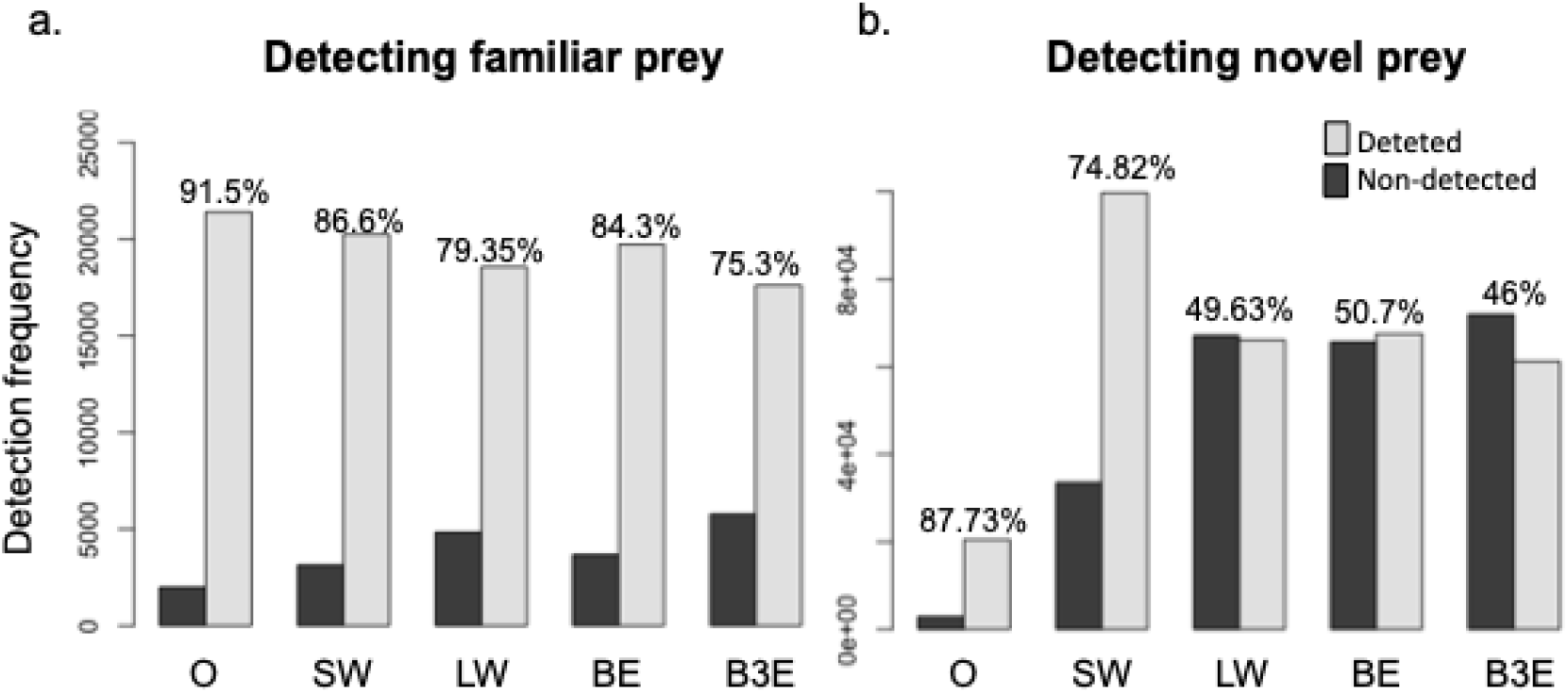
Detection frequency per DNN and morph at detecting familiar prey (a, training DNNs on all prey forms and the background) and novel prey (b, training DNNs on only background and the opaque form). Results for all DNN architectures are pooled together. Percentage of detection per morph are represented on top of the bars.

### Comparisons between birds and DNNs

For both types of observers, the O morph was among the easiest morphs to detect, while morphs touching 3 edges were among the more difficult morphs to detect. However, differences in the detectability of different morphs were not similar between field experiments and DNNs detecting either familiar or novel prey (Table S4 for Mantel tests comparing field experiments and all DNNs mean score per morph). When prey forms are pooled according to window surface area or the number of broken edges, field work experiments and DNNs also produce different results (Tables S4 and S5).

## Discussion

### Important features of transparent elements

By studying prey detection by natural avian predators, we found that in addition to their presence, the position of transparent elements is key at reducing prey detectability. Moths with transparent elements touching wing edges were more efficient at reducing detectability in our field experiment. Our results are consistent with the findings of [32] that breaking prey contour is more efficient at reducing prey detectability than 1) having a uniform colour that matches one of the several colours present in the background, and 2) sporting elements that represent a random sample of the background, which is similar to the visual effect created by large transparent elements. Surprisingly, however, the disruption of wing outline does not seem to be the dominant strategy used by butterflies and moths exhibiting transparent wing elements. Several clearwing Lepidoptera have well defined wing borders, suggesting that factors other than reducing detection by predators are also shaping transparency evolution. Transparency in wings is sometimes restricted to small areas, and/or is combined with highly visible elements, suggesting an important role in visual communication, beyond crypsis, as in several unpalatable comimics of the Ithomiinae tribe [33]. It is also possible that in Lepidoptera, evolving large transparent areas touching wing edges is constrained by factors linked to wing strength or flight dynamics. Whether transparent zones without borders are indeed more fragile or sensitive to wind and/or water remains an open question.

### Can DNNs predict avian predation?

Both natural predators and DNNs detected “O” the most easily, whereas “B3E” was among the hardest morphs to detect for both observers. This suggests that transparent elements breaking several prey edges should indeed be favoured as they are more difficult for predators to detect, a benefit that could ease their establishment in a cryptic opaque population. Our results also suggest that DNNs can be used to predict predators’ behaviour to a certain extent. However, their behaviours toward the other morphs were rather different during our study. Differences between morph detection is small for both types of observers, although these differences are often significant for DNNs. Grouping morphs by their characteristics also did not produce similar results between wild predators and DNNs. While broken edges reduced predation rates in the field experiment, DNNs detected a stronger effect of size rather than position of the window on decreasing detection. This difference was especially high when DNNs detected novel prey, which was the simulation expected to be closer to field experiments as the local community of predators are expected to be familiar with opaque and not with transparent prey.

One potential explanation for the lack of agreement between the field and DNN results could be related to what we were actually testing in our experiments. The DNN experiment only explored the detectability of different prey items. By contrast, the field experiment not only tested the detectability of different prey items, but also the decision of the predator to attack, without the possibility to disentangle which of these factors was more important. Both detection and motivation to attack in biological neural networks are likely mediated by both top-down (e.g. neophobia, hunger, attention) and bottom-up (feature-driven) cognitive processes, whereas the DNNs we used make use of bottom-up processes only. This may partly explain why DNNs and birds did not show the same relative ability to detect the different morphs.

Another possibility is that the attack rate in the field experiment was too high to see differences in predation rates among morphs. The attack rate in this experiment was over double that of a similar study carried out under similar conditions (35.1% in the current study vs 14.08% in [4]). Under strong predation pressure, once the most detectable prey morphs in a given area have already been attacked, less detectable prey are more likely to be attacked. This could potentially have led to a homogenisation of attack rates among morphs. Another possibility is that there was some kind of unintentional bias in the placement of moths in the field experiment. While in the DNN experiments all DNNs had access to images in which we were sure the moths were visible, this may not have been the case for all prey items in the field experiment. Natural predators may have mostly attacked the moths placed in the most visible locations. If visible locations unintentionally contained more of certain morphs, then these morphs may have been more heavily predated upon than would otherwise be expected.

Other potential explanations for the difference between the field and DNN experiments could be related to differences in the colour and achromatic contrasts seen by birds and DNNs. DNNs were fed standard RGB photos designed to mimic the colours experienced by human observers. Avian observers possess a UV cone and three visible-light cones that are more spectrally separated than the three visible-light cones of humans, indicating that birds may see greater colour contrasts than humans. That said, birds experience higher receptor noise than humans do in both their achromatic and their colour vision channels [34]. The greater noise birds experience in their colour vision channel must, to some extent, counteract their greater number of more spectrally separated cones. Automatic image processing software on board the iPad and the smartphone that took the pictures can be expected to have artificially enhanced both achromatic and colour contrasts beyond what would be seen by human observers. Achromatic contrasts were therefore likely exaggerated beyond what an avian observer would perceive. Colour contrasts must also have differed, but, for the reasons mentioned above, it is not straightforward to predict in which direction.

The spatial configuration of elements seen by birds and DNNs may have also differed. Birds may search for moths from a different viewing angle than the moths were photographed from. All photographs were taken from a viewing angle perpendicular to the surface of the moth wings. However, birds may search while flying or walking on the trunk or the ground. Different viewing angles can be expected to impact not only the apparent configuration of opaque elements, but also the specular reflection (i.e. glare) by transparent elements. In the first set of images in Figure 4, some specular reflection can be seen from the transparent windows of the BE morph. Real butterflies and moths with transparent elements show comparatively less specular reflection; thus, it would be interesting to run a similar set of experiments with artificial moths containing empty windows rather than transparent windows made from synthetic materials.

It is, of course, possible that the lack of agreement between live predators and DNNs could be due to limitations of the DNNs themselves. Although DNNs have made incredible strides in mimicking biological perception in the last decade, they are still susceptible to error in ways that computer scientists are still discovering [35]. It was not practical to analyse all 363,000 class activation maps (CAMs) produced by our dataset, but we did notice a strong bias towards the highest activations overlapping the anterior wing edges, with a slight bias toward the left anterior edge, regardless of whether the DNN was trained on the opaque or on all morphs, and regardless of whether the images were mirrored such that the left side became the right side and vice versa. The importance of diagonally-oriented wing edges is particularly evident in the bottom row of images in Figure 4, where moderate activations were produced in response to a diagonally-oriented tree branch viewed against the sky. It is interesting to note that the anterior wing edges of the BE morph were thicker than those of the LW morph. If the anterior wing edge was more important than the posterior wing edge for correct categorization by DNNs, but not for live predators, this may explain why DNNs detected the BE morph more easily than the LW morph, but the opposite was true of live birds.

Recent studies have shown that DNNs do not encode global object shape, but rather local object features, and perform poorly when asked to classify images composed of silhouettes without any internal object texture [36]. Indeed, Baker et al. [37] found VGG-19’s performance to be abysmal when objects had clearly-defined shapes against a white background but incongruous internal textures. This represents a key difference between human and DNN perception and is perhaps the most likely explanation for why the DNNs’ and avian predators’ responses did not mirror each other. Moving forward, we would advise caution in assuming that DNNs will mimic the finer points of biological perception in simulations like ours without validating that this is true in live animals. However, we remain optimistic that studies like this one may lend some insight to computer scientists attempting to develop DNN algorithms that model biological perception more accurately.

## Conclusions

According to both the DNN and fieldwork experiments’ results, having transparent elements breaking several prey borders seems to be the most effective mechanism for reducing detectability of prey and subsequent predation. However, to be able to directly compare DNNs’ and wild predators’ reactions, careful experimental design, paying special attention to what may affect their output, is needed.

## Supporting information

ESM

## Acknowledgements

This work was funded by Clearwing ANR project (ANR-16-CE02-0012), HFSP project on transparency (RGP0014/2016) and a France-Berkeley fund grant (FBF #2015--58). With the support of LabEx CeMEB, an ANR “Investissements d’avenir” program (ANR-10-LABX-04-01).

