## Supplementary material for "Deep neural network and field experiments reveal how transparent wing windows reduce detectability in moths": ESM

**Electronic supplementary material**

Affiliations:

*first co-authors

**Extended Material and Methods**

Field experiments

We followed an experimental design similar to that described in [1]. We performed predation experiments in April and May 2019 in two localities in southern France: La Rouvière forest (43.65°N, 3.64°E) and the Montpellier zoo (43.64°N, 3.87°E), for three 1-week sessions at each place. We monitored artificial prey survival from predation by bird communities that include great tits (*Parus major*), blue tits (*Cyanistes caeruleus*), Eurasian jays (*Garrulus glandarius*)*,* common chaffinches (*Fringilla coelebs*), golden orioles (*Oriolus oriolus*) and European robins (*Erithacus rubecula*). Artificial prey (body and wings) were pinned on green oak (*Quercus ilex*) tree trunks (>10cm in diameter, with little or no moss cover) every 10m. To deter ant attacks, we applied Vaseline and double-sided transparent tape between the wings and the trunk. Prey were placed perpendicularly to the ground and mostly on the north-facing side of tree trunks to reduce direct sunlight reflection that could facilitate their detection. We monitored prey survival once per day for the following four consecutive days after placing them on trunks, and removed them afterwards.

Artificial moths

As described by Arias *et al* [1] and similarly to other experiments, artificial moths consisted of paper wings and an edible body [2,3]. Wings consisted of right triangles resembling resting generic moths (i.e., not representing any real species). Triangle dimensions were 25mm height by 36mm width, thus a surface area of 450mm². To determine wing colouration, we performed reflectance measurements of green oak trunk colouration (120 measurements on 6 trunks) and laminated grey paper using a deuterium halogen lamp (Avalight DHS) emitting over 300-700 nm range including UV to which birds are sensitive [4], a spectrophotometer (Starline Avaspec-2048 L), an optic probe with tip cut at 45° (FC-UV200-2-1.5 x 100, Avantes), and a white diffuse reference (spectralon, WS2). We calculated colour and brightness contrasts between paper and trunk as seen by birds by applying Vorobyev and Osorio discriminability model [5] using *pavo* package with R software [6]. We modeled the discriminality for both UVS vision (blue tit, [7]) and VS vision (shearwater, [8]). We used a forest shade light environment [9], and calculated Weber fractions for the chromatic response per cone type by dividing the standard deviation of noise in a single cone, i.e. 0.1 (mean calculated from Lind et al. [10] and Olsson et al. [11]), by the square root of the number of cones per neurally integrative unit [5]. These cone numbers were estimated as Order-level means of cone ratios of terrestrially foraging birds to be 1:1.7:2.5:3 (U/V:S:M:L) [8,12–15]. A Weber fraction of 0.2 was used for the brightness response (as the average reported values for known bird species [16]). We found that colour Grey155 (R=G=B=155), printed on Canson® sketch paper with a HP officejet pro 6230 printer was chromatically indistinguishable from trunk coloration (chromatic contrast of 0.47±0.16 JND for UVS vision and of 0.41±0.14 JND for VS vision), and marginally lighter than oak trunks (achromatic contrast of 1.65±0.69 JND for UVS vision and of 1.65±0.68 JND for VS vision). This colour thus enabled us to explore the effect of transparent windows and their spatial configuration on reducing detectability of cryptic prey.

Data analysys: effect of local bird demography on attack rate

Differences in attack rates between places and/or weeks could be produced by differences in predation pressure, which can be higher when more nests are occupied and/or when more chicks at their developmental peak are present in the population. Therefore, we explored whether differences in attack rate per week/locality were associated with great and blue tit chick hatching, the number of chicks between 9 and 15 days old (age when the food demand by great tit and blue tit chicks is the highest (A. Charmantier pers. comm.)), or the number of active nests or number of nests having chicks between 9 and 15 days old. We fitted linear models with attacks per day as the response variable and including all, some or none of these demographic variables as explanatory variables. We selected the best model using the AIC criterion.

Deep Neural Networks

Initial learning rate and solvers

The initial learning rate in subsequent layers was set to 2*10^-4^, and was increased by a factor of ten in the final, fully-connected layer. The L_2_ regularization term was set to 0.0001. Different solvers, which identify optimal weights, were used for different DNNs to maximize the learning capability (measured as the test set accuracy) of the different DNN architectures. The adaptive moment estimation (Adam) solver was used for AlexNet, ResNet-18, SqueezeNet, and GoogLeNet, and the stochastic gradient descent with momentum (sgdm) solver was used for VGG-16 and VGG-19.

Training methods

We used two training methods to explore two different questions: 1) In a community of different moth morphs, all of which are familiar to predators, which morphs will be the most successful at evading predation? and, 2) If predators are only experienced with opaque morphs, which novel morph(s) with transparent elements will most easily establish themselves in the population? (Fig. 3).

*Detection of familiar morphs by DNNs*

We trained DNNs to detect moths using all morphs at once to test the effect of transparent elements at reducing detectability among familiar prey. Out of the 222 images of each morph, the training set included 144 randomly-selected images of each of the five morphs and each corresponding background image in quintuplicate. We included each background in quintuplicate so that there would be the same number of positive (with a moth) and negative stimuli (without a moth) in the training set. Thus, in total, 1,440 images were used in training. Thirty-nine randomly-selected images of each morph and their corresponding background images in quintuplicate were used in the validation set (for a total of 390 images), and the remaining 39 images (out of 222) of each morph and the background were used in the test set (for a total of 234 images). We repeated this training and testing procedure 100 times, each time retraining the same initial DNN (pretrained on the ImageNet database) and randomly selecting which images (out of the full 222 images available per morph) were to be allocated for training, for validation, and for testing. Minibatch size (the subset of images used during one iteration of training) was set to 80 for AlexNet, ResNet-18, SqueezeNet, and GoogLeNet, and to 70 for VGG-16 and VGG-19 (lower in these latter two due to computer memory limitations). Each minibatch contained eight (for AlexNet, ResNet-18, SqueezeNet, and GoogLeNet) or seven (for VGG-16 and VGG-19) sets of ten images. Each set of ten images had the same background, five of which contained one each of the O, SW, LW, BE, and B3E morphs. The remaining five in each set of ten was the same background image in quintuplicate. Minibatch sizes during training were larger for this than for the training method described below to keep the number of different backgrounds in each minibatch the same under each training scenario.

*Which new morph would establish in a population?*

In the second experiment, we trained DNNs to discriminate between pictures with a moth present on a trunk (O morph only) and pictures of the background only. The aim was to simulate the reaction of predators familiar with opaque prey and naive to prey exhibiting transparent elements. This approach allowed us to test which new visual configuration is less often detected, and thus potentially more likely to establish in a population. Each time, the training set of images consisted of 144 randomly-selected images of the O morph and 144 images of the corresponding backgrounds (with no moth present), the validation set included 39 randomly-selected images of the O morph and 39 images of the corresponding background, and the test set consisted of the remaining 39 (out of 222) images of the O morph, 39 images of the corresponding background and 222 images of each of the other morphs. As before, in each of these sets of images, the positive and negative stimuli had the same visual backgrounds; only the morph type varied. The minibatch size was set to 16, and each minibatch contained eight O morph images and eight background images that had the same backgrounds as the O morph images. All other training parameters were identical to those above.

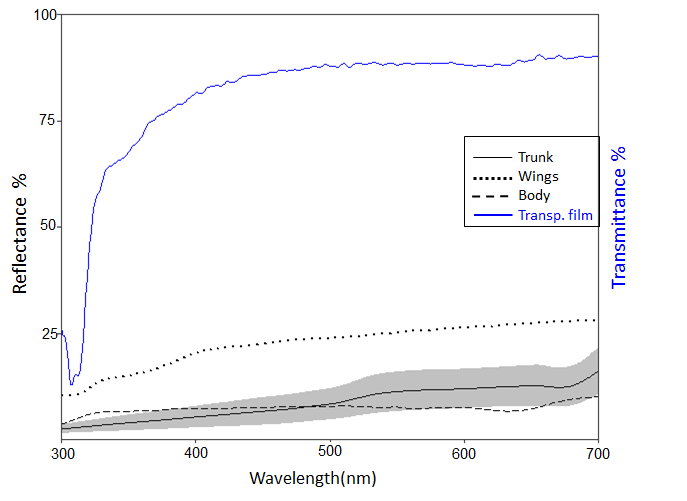

Figure S1. Reflectance spectra of the background (oak tree trunks in solid black line with a confidence interval of ± 1 standard deviation) and opaque areas: body (dashed line) and opaque wings (dotted line), and transmittance of the transparent film (solid blue line).

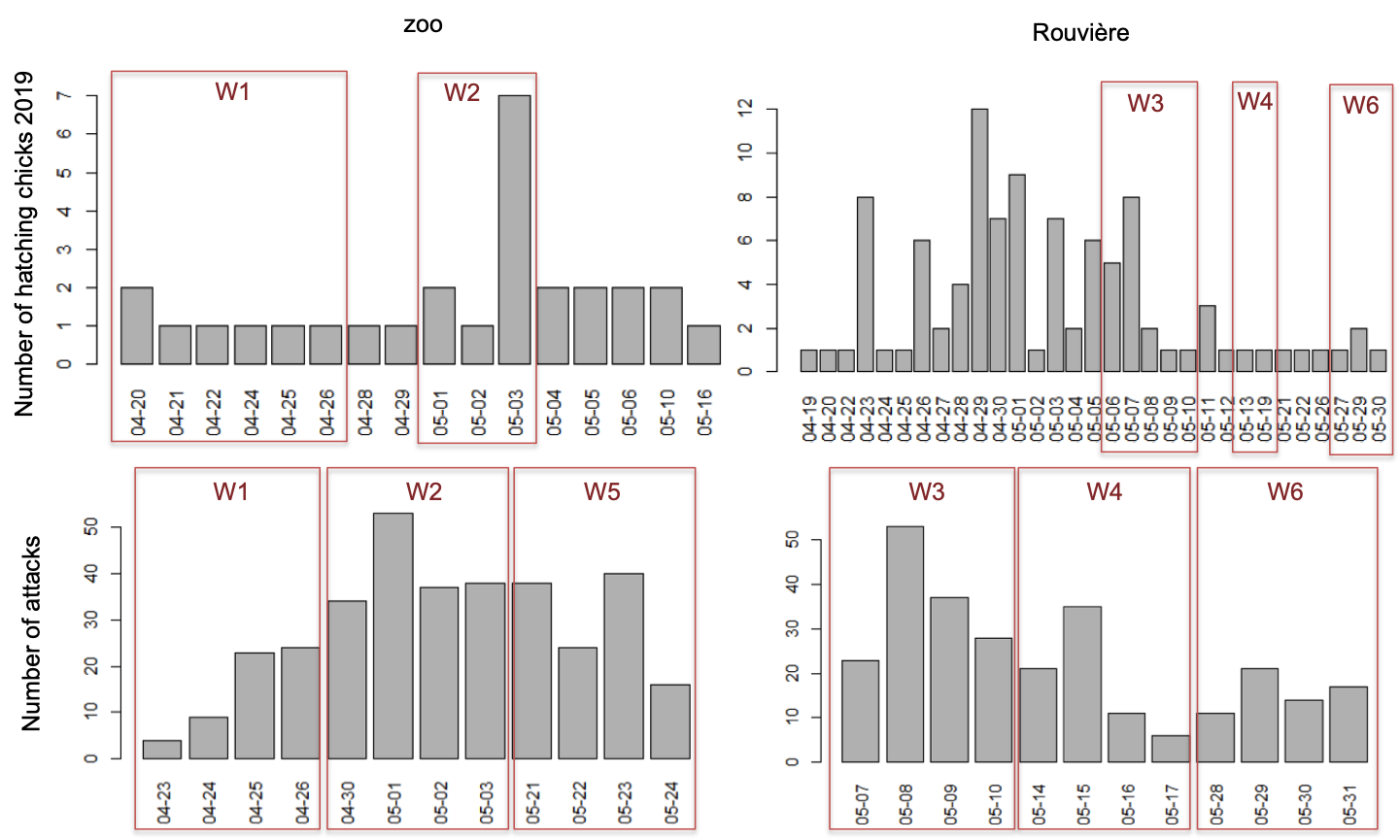

Figure S2. Hatching chicks (top) and attacked prey (bottom) at the two different experimental sites: the zoo (left) and La Rouvière forest (right). Red squares in the top plots indicate the days when the experiment was run in each of the places and in the bottom plot delimit the weeks of the experiment. By week 5 no more hatchlings were registered at the zoo during 2019.

Table S1. False positive rate for different DNNs and training protocols

| Training morph | DNN | False positive rate |
| --- | --- | --- |
| O | ResNet | 0.01 |
| O | SqueezNet | 0.06 |
| O | AlexNet | 0.19 |
| O | GoogLeNet | 0.03 |
| O | vgg16 | 0.03 |
| O | vgg19 | 0.03 |
| all | SqueezNet | 0.18 |
| all | ResNet | 0.12 |
| all | AlexNet | 0.31 |
| all | vgg16 | 0.12 |
| all | vgg19 | 0.14 |
| all | GoogLeNet | 0.09 |

Table S2. Summary statistics on moth detection per DNN architecture and moth morph when DNN detected familiar (trained in all morphs, left) and novel prey (trained on opaque, right).

| DNN | Morph | Detection of familiar prey (mean±SD) | Detection of novel prey (mean±SD) |
| --- | --- | --- | --- |
| AlexNet | O | 0.81±0.39 | 0.75±0.43 |
| AlexNet | SW | 0.72±0.45 | 0.61±0.49 |
| AlexNet | LW | 0.61±0.49 | 0.47±0.5 |
| AlexNet | BE | 0.71±0.45 | 0.55±0.5 |
| AlexNet | B3E | 0.62±0.48 | 0.47±0.5 |
| GoogLeNet | O | 0.93±0.25 | 0.91±0.29 |
| GoogLeNet | SW | 0.88±0.32 | 0.78±0.41 |
| GoogLeNet | LW | 0.81±0.39 | 0.54±0.5 |
| GoogLeNet | BE | 0.86±0.35 | 0.54±0.5 |
| GoogLeNet | B3E | 0.76±0.43 | 0.5±0.5 |
| ResNet | O | 0.93±0.25 | 0.87±0.34 |
| ResNet | SW | 0.9±0.3 | 0.75±0.43 |
| ResNet | LW | 0.85±0.36 | 0.47±0.5 |
| ResNet | BE | 0.88±0.33 | 0.44±0.5 |
| ResNet | B3E | 0.8±0.4 | 0.41±0.49 |
| SqueezNet | O | 0.93±0.26 | 0.9±0.3 |
| SqueezNet | SW | 0.87±0.33 | 0.73±0.44 |
| SqueezNet | LW | 0.8±0.4 | 0.5±0.5 |
| SqueezNet | BE | 0.86±0.34 | 0.53±0.5 |
| SqueezNet | B3E | 0.79±0.41 | 0.49±0.5 |
| vgg16 | O | 0.94±0.24 | 0.92±0.28 |
| vgg16 | SW | 0.9±0.3 | 0.81±0.39 |
| vgg16 | LW | 0.83±0.37 | 0.52±0.5 |
| vgg16 | BE | 0.87±0.34 | 0.52±0.5 |
| vgg16 | B3E | 0.76±0.43 | 0.45±0.5 |
| vgg19 | O | 0.95±0.23 | 0.92±0.27 |
| vgg19 | SW | 0.91±0.28 | 0.8±0.4 |
| vgg19 | LW | 0.85±0.36 | 0.48±0.5 |
| vgg19 | BE | 0.88±0.32 | 0.46±0.5 |
| vgg19 | B3E | 0.79±0.41 | 0.44±0.5 |

Table S3. Generalised linear mixed model results for average morph detection for all DNNs in both experiments: detection of familiar prey (left, DNN trained on all morphs and background) and detection of novel prey (right, DNN trained on O morph and background).

|  | Detection of familiar prey | | | Detection of novel prey | | |
| --- | --- | --- | --- | --- | --- | --- |
|  | Estimate±SE | z | P | Estimate±SE | z | P |
| (Intercept) | 4.38±0.42 | 10.36 | *** | 1.75±0.15 | 11.7 | *** |
| O >SW, LW, BE, B3E | 0.41±0.04 | 9.8 | *** | 0.76±0.04 | 20.25 | *** |
| SW>LW, BE, B3E | 0.29±0.06 | 4.66 | *** | 0.6±0.06 | 10.87 | *** |
| BE > B3E | 0.68±0.12 | 5.67 | *** | 0.26±0.11 | 2.39 | * |
| full windows > broken windows | -0.03±0.1 | -0.32 |  | 0.08±0.09 | 0.85 |  |

Prey morphs are the explanatory variables (O: opaque, SW: small windows, LW: large windows, BE: windows touching one border, B3E: windows touching three borders). Symbols: . * p < 0.05, *** p < 0.001.

Table S4. Mantel test results between distance matrices of field experiments and all DNNs average mean score per morph when detecting familiar (trained in all morphs) and novel prey (trained on opaque).

|  | Comparisons | Observation (R) | Simulated p |
| --- | --- | --- | --- |
|  | Field vs mean DNN (opaque) | 0.106 | 0.23 |
| Per morph | Field vs mean DNN (all) | 0.081 | 0.42 |
|  | mean DNN (all) vs mean DNN (opaque) | 0.71 | 0.01 |
|  | Field vs mean DNN (opaque) | 0.939 | 0.17 |
| Per surface | Field vs mean DNN (all) | 0.828 | 0.17 |
|  | mean DNN (opaque) vs mean DNN (all) | 0.97 | 0.16 |
|  | Field vs mean DNN (opaque) | 0.99 | 0.16 |
| Per border | Field vs mean DNN (all) | 0.1 | 0.5 |
|  | mean DNN (opaque) vs mean DNN (all) | -0.005 | 0.51 |

Tables S5. Distance matrices per morph, border and surface for field, detection of familiar prey and detection of novel prey experiments. For the field experiment they are calculated from the Cox survival coefficients. For the DNNs experiments they are calculated from the average score for all DNNs and replicates. They are always calculated by subtracting values corresponding to column morphs - values corresponding to row morphs.

Tables S5.1. Per morph

Table S5.1.a From the field

| Field | O | SW | LG | B1E | B3E |
| --- | --- | --- | --- | --- | --- |
| O | 0 | -0.02 | -0.02 | -0.16 | -0.22 |
| SW | 0.02 | 0 | 0 | -0.14 | -0.2 |
| LG | 0.02 | 0 | 0 | -0.14 | -0.2 |
| BE | 0.16 | 0.14 | 0.14 | 0 | -0.06 |
| B3E | 0.22 | 0.2 | 0.2 | 0.06 | 0 |

Table S5.1.b From detection familiar prey experiment

| All | O | SW | LG | BE | B3E |
| --- | --- | --- | --- | --- | --- |
| O | 0 | -0.05 | -0.12 | -0.07 | -0.16 |
| SW | 0.05 | 0 | -0.07 | -0.02 | -0.11 |
| LG | 0.12 | 0.07 | 0 | 0.01 | -0.04 |
| BE | 0.07 | 0.02 | -0.01 | 0 | -0.09 |
| B3E | 0.16 | 0.11 | 0.04 | 0.09 | 0 |

Table S5.1.c. From detection novel prey experiment

| Opaque | O | SW | LG | BE | B3E |
| --- | --- | --- | --- | --- | --- |
| O | 0 | -0.13 | -0.38 | -0.37 | -0.42 |
| SW | 0.13 | 0 | -0.25 | -0.24 | -0.29 |
| LG | 0.38 | 0.25 | 0 | 0.01 | -0.04 |
| BE | 0.37 | 0.24 | -0.01 | 0 | -0.05 |
| B3E | 0.42 | 0.29 | 0.04 | 0.05 | 0 |

Tables S5.2. Per surface

Table S5.2.a From field experiment

| Field | O | SW | LargeW |
| --- | --- | --- | --- |
| O | 0 | -0.02 | -0.13 |
| SW | 0.02 | 0 | -0.11 |
| LargeW | 0.13 | 0.11 | 0 |

Table S5.2.b. From detection familiar prey experiment

| All | O | SW | LargeW |
| --- | --- | --- | --- |
| O | 0 | -0.05 | -0.12 |
| SW | 0.05 | 0 | -0.07 |
| LargeW | 0.12 | 0.07 | 0 |

Table S5.2.c. From detection novel prey experiment

| Opaque | O | SW | LargeW |
| --- | --- | --- | --- |
| O | 0 | -0.13 | -0.39 |
| SW | 0.13 | 0 | -0.26 |
| LargeW | 0.39 | 0.26 | 0 |

Table S5.3. Per border

Table S5.3.a From field experiment

| Field | Complete | BE | B3E |
| --- | --- | --- | --- |
| Complete | 0 | -0.15 | -0.21 |
| BE | 0.15 | 0 | -0.06 |
| B3E | 0.21 | 0.06 | 0 |

Table S5.3.b. From detection familiar prey experiment

| All | Complete | BE | B3E |
| --- | --- | --- | --- |
| Complete | 0 | -0.02 | -0.11 |
| BE | 0.02 | 0 | -0.09 |
| B3E | 0.11 | 0.09 | 0 |

Table S5.3.c. From detection novel prey experiment

| Opaque | Complete | BE | B3E |
| --- | --- | --- | --- |
| Complete | 0 | -0.14 | -0.18 |
| BE | 0.14 | 0 | -0.05 |
| B3E | 0.18 | 0.05 | 0 |
